# A novel unbiased test for molecular convergent evolution and discoveries in echolocating, aquatic and high-altitude mammals

**DOI:** 10.1101/170985

**Authors:** Amir Marcovitz, Yatish Turakhia, Michael Gloudemans, Benjamin A. Braun, Heidi I. Chen, Gill Bejerano

**Affiliations:** Department of Developmental Biology, Stanford University, Stanford, California, 94305, USA; Department of Electrical Engineering, Stanford University, Stanford, California, 94305, USA; Biomedical Informatics Program, Stanford University, Stanford, CA 94305, USA; Department of Computer Science, Stanford University, Stanford, California 94305, USA; Department of Pediatrics, Stanford University, Stanford, California 94305, USA

**Keywords:** convergent evolution, genome wide tests, echolocation, aquatic, high altitude, subterranean, mammals

## Abstract

Distantly related species entering similar biological niches often adapt by evolving similar morphological and physiological characters. The extent to which genomic molecular convergence, and the extent to which coding mutations underlie this convergent phenotypic evolution remain unknown. Using a novel test, we ask which group of functionally coherent genes is most affected by convergent amino acid substitutions between phenotypically convergent lineages. This most affected sets reveals 75 novel coding convergences in important genes that pattern a highly adapted organ: the cochlea, skin and lung in echolocating, aquatic and high-altitude mammals, respectively. Our test explicitly requires the enriched converged term to not be simultaneously enriched for divergent mutations, and correctly dismisses relaxation-based signals, such as those produced by vision genes in subterranean mammals. This novel test can be readily applied to birds, fish, flies, worms etc., to discover more of the fascinating contribution of protein coding convergence to phenotype convergence.

## Introduction

The evolutionary gain of similar traits in distantly related species has been proposed to be encoded, in some cases, by identical mutational paths in their respective genomes^1,2^. While convergent phenotypes are seldom, if ever, the result of only convergent molecular adaptations, several cases of identical amino acid changes promoting phenotypic convergence in distant species have been documented in recent years^3^. For example, parallel evolution of *Prestin* (*SLC26A5*)^4–6^ and several other auditory genes^7^ was identified in echolocating bats and toothed whales. However, known examples are few and relatively limited in their biological scope. The opportunity to explore the genetic basis of a diverse array of natural adaptations warrants more systematic approaches for identifying adaptive molecular convergence patterns across genomes^8,9^. Recent genome-wide screens illuminated sensory genes with sequence changes that segregate with echolocating mammals^8^, and others highlighted positively selected genes that contain convergent amino acid substitutions in independent lineages of aquatic mammals^9,10^. However, later analyses demonstrated that the genome-wide frequency of molecular convergence in echolocating mammals is similar to the frequency in non-echolocating control outgroups, rendering the true magnitude of the phenomena and the significance of molecular convergence largely unknown^11,12^.

Another key challenge in determining the molecular basis of phenotypic convergence is distinguishing adaptive convergent mutations from sets of mutations with molecular convergent-like patterns that do not contribute to the phenotypic convergence, but instead accumulate for other reasons^13^. In particular, relaxation of evolutionary constraints^14,15^ or complete loss of a biological function^16^ may lead to an accumulation of convergent-like (but in fact non-adaptive) mutations in functionally related groups of genes.

Here we develop an unbiased approach to determine whether we can detect a molecular signature of phenotypic convergence in two distantly related lineages by observing adaptive coding convergence in their genomes. Scanning the genomes of multiple convergent mammalian species (Fig. 1a) for amino acid convergence events (Fig. 1b), we devised a novel, functional enrichment-based analysis, that rather than ask which individual genes contain convergent amino acid mutations, asks which biological function, if any, is most significantly affected by coding convergence. The test is unbiased because it collects convergent mutations from all genes, and because it is willing to declare any one of thousands of different functions as its top enriched one. We also make the test specifically robust to constraint relaxation by detecting concurrent accumulation of divergent mutations^12,13^ (Fig. 1c) in these same genes. The novel method is successfully applied to three convergent and three relaxation-based scenarios to discover fascinating molecular convergence events in highly convergent organs.

**Figure 1.**
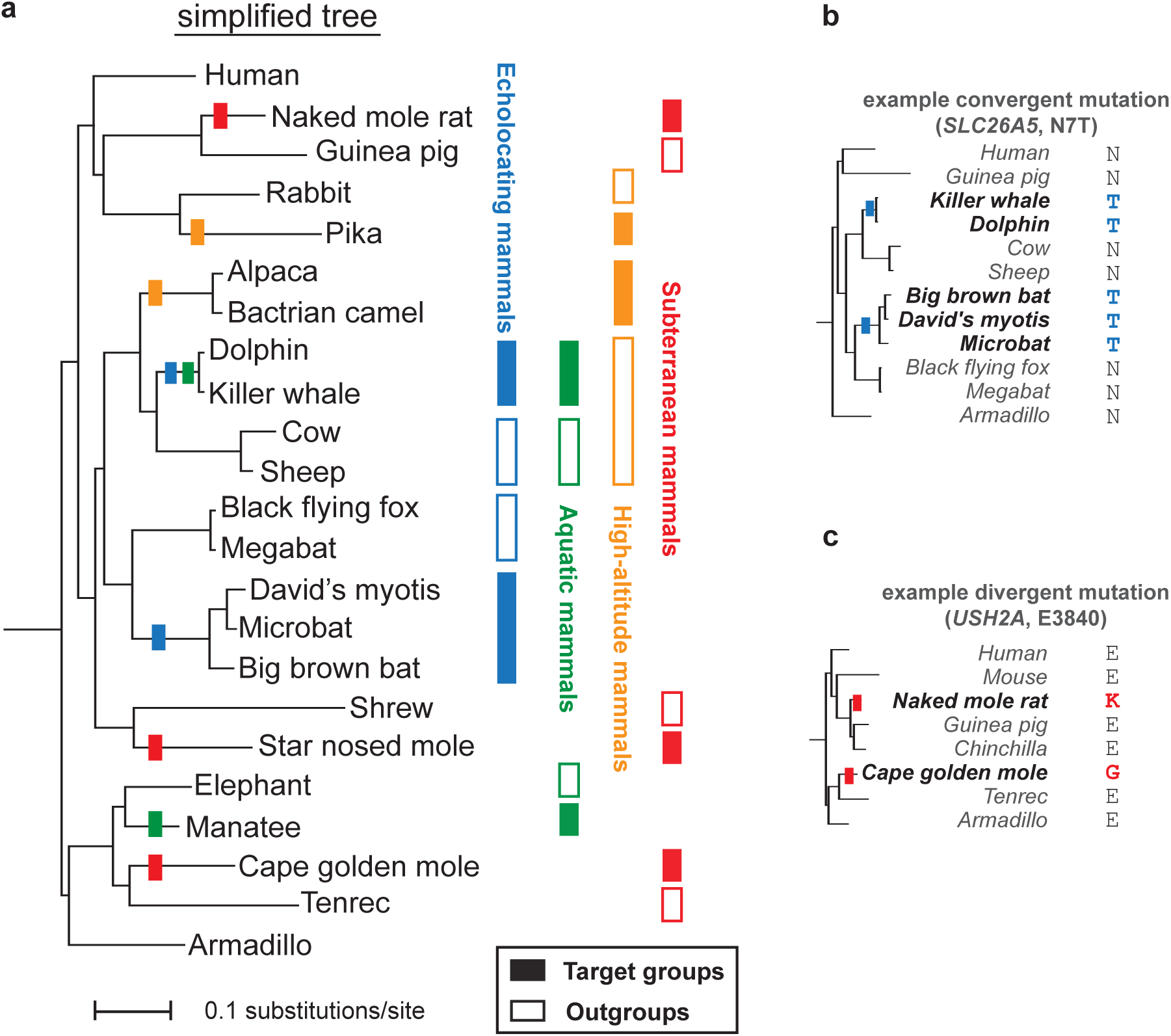
Searching for molecular convergence and divergence in the mammalian lineage. **a,** Left: reduced placental mammals phylogenetic tree. See Supplementary Fig. 1 for all 57 species used. Colored rectangles highlight branches with independent phenotypic evolution of echolocation (blue), aquatic (green), high-altitude (orange), and subterranean (red) lifestyles. Right: Filled and empty rectangles represent target and outgroup species, respectively. We screen for mutations along the branches leading from the last common ancestor of each target group and its outgroup to the target group itself. For example: **b**, An N7T (Asparagine to Threonine in position 7) convergent mutation in the hearing gene prestin (*SLC26A5*) in echolocating mammals. **c**, An E3840 divergent mutation in the vision/hearing gene usherin (*USH2A*) in a pair of distantly related subterranean mammals.

## Results

### Convergent amino acid substitutions in echolocating bats and cetaceans point to the cochlear ganglion

Toothed whales (*Odontoceti*) and certain bat lineages have independently evolved a sophisticated sonar system for three-dimensional orientation and prey localization in conditions where vision is ineffective, such as at night or in turbid water^17^. Starting with 57 placental mammalian whole genomes available at UCSC (Supplementary Table 1), we first selected two echolocating species groups (dolphin and killer whale; David’s myotis, microbat and big brown bat) and two respective outgroups (cloven-hoofed mammals and megabats; Fig. 1a and Supplementary Figure 1). We then looked for all human protein coding genes that: have an ortholog in at least 40 of the 56 other mammals, have at least one amino acid highly conserved across the orthologs, and are annotated for specific function using the MGI phenotype ontology^18^. This identifies 2,988,146 conserved (and thus likely functionally important) amino acids across 6,719 genes annotated with 4,300 different MGI functions (Fig. 2a and Methods). Scanning all conserved amino acids for convergence events between the bats and the whales, placed exactly on the branch separating them from their outgroup (see Methods and Fig. 1b) reveals 770 such events across 656 genes (Supplementary Tables 2 and 3). We also produced three control sets, where one or both outgroup species were swapped with their respective target group. The three control pairing combinations tested a very similar number of positions (2,986,063-2,987,561) and produced a very similar number of convergence events, involving 701–770 amino acids in 593-658 genes, confirming that no excess genome-wide convergence exists between echolocating cetaceans and bats relative to other pairs^11,12^ (Fig. 2b).

**Figure 2.**
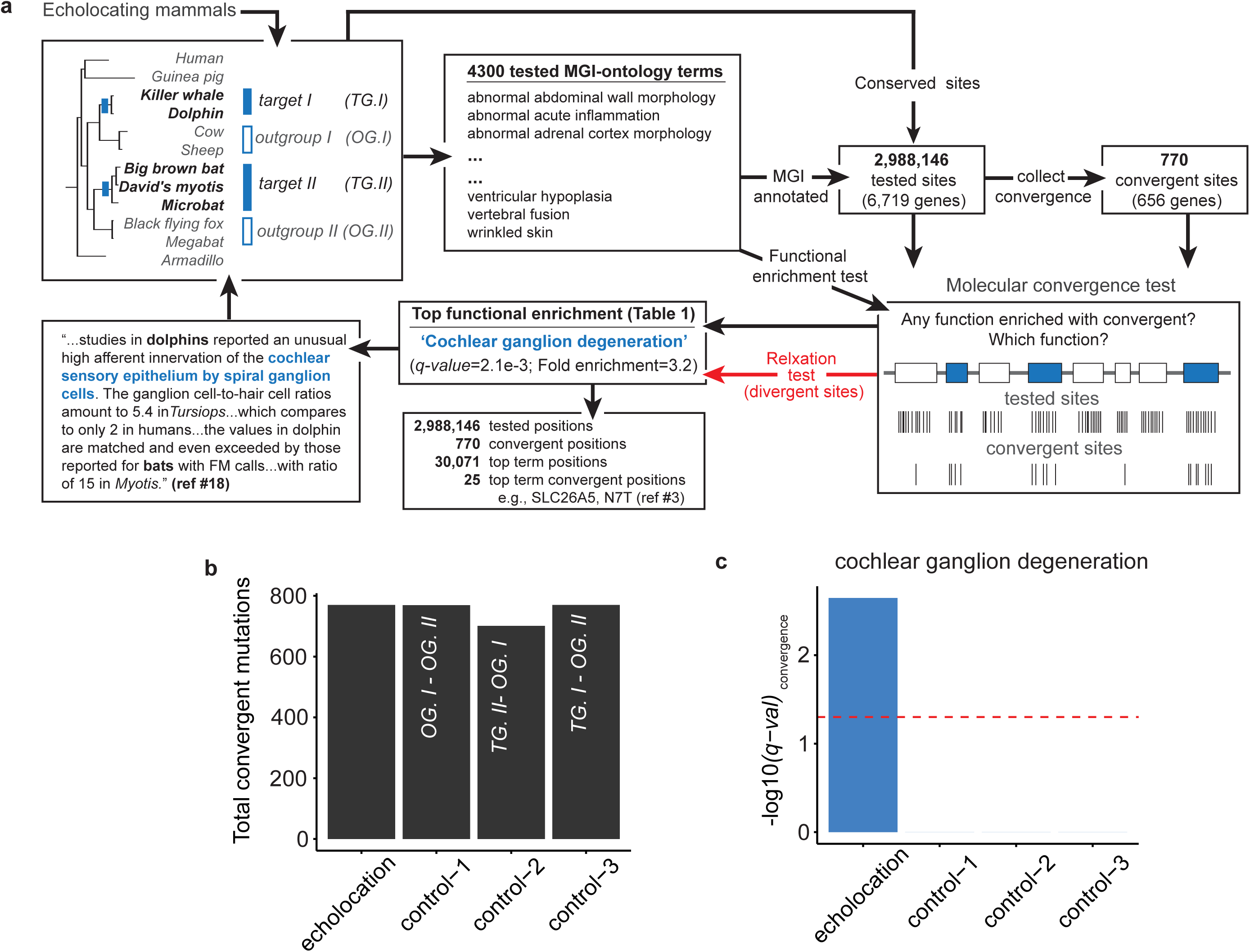
A novel molecular convergence test. Example application to echolocating mammals: **a**, Top left. We pick 2 target groups (TG) of species with a phenotypic convergence (echolocation here), and two outgroups (OG). Top middle. Our algorithm identifies all cross-species conserved (and thus functionally important) amino acid positions in genes annotated for any of 4,300 specific MGI phenotype functions. Top/bottom right. We then identify a subset of positions showing convergent mutations between our target groups, and perform a hypergeometric test over positions to find the single most statistically enriched MGI function (if any) for amino acid convergence. We also test for divergent mutations between our target groups (Red). Bottom middle/left. If the most enriched convergent function is *not* also enriched for divergent mutations we declare an adaptive molecular convergence event, and link it back to the convergent species phenotypes. In this example we discover 25 convergence events in 18 important hearing genes in bats and whales, prime fodder for concerted follow-up work. The cochlear ganglion prediction is particularly striking considering that 4,300 different functions were evaluated. **b**, A comparable total number of convergent mutations is observed in echolocating mammals (TG. *I* and *II* in panel a top left), and three control sets formed by shuffling either or both target groups with their cognate outgroup. **c**, However, only the echolocating (left) set in panel b shows a statistically significant enrichment for molecular convergence (of the cochlear ganglion). None of the control groups shows any statistically enriched term.

We then asked whether each converged set of amino acids was enriched for any particular MGI phenotype term, by computing its q-value, a multiple hypothesis corrected p-value (see Methods). Testing and correcting for 4,300 MGI terms, the single most enriched function for convergent mutations in echolocating mammals is (genes whose mutations have been observed to cause) **‘cochlear ganglion degeneration’** (*q-value*=2.4e-3, hypergeometric test; Table 1), with 25 convergent mutations in 18 genes (Fig. 2a, Supplementary Table 3). The cochlear ganglion is the group of nerve cells that serve the sense of hearing by sending a representation of sound from the cochlea to the brain. Indeed, both dolphins and microbats display a significant increase in ganglion to hair cell ratio in their inner ear cochleas compared to non-echolocating mammals^19^. In contrast, there were no statistically significant functional enrichments for molecular convergence in the three sets of control target groups (Fig. 2c).

**Table 1.**
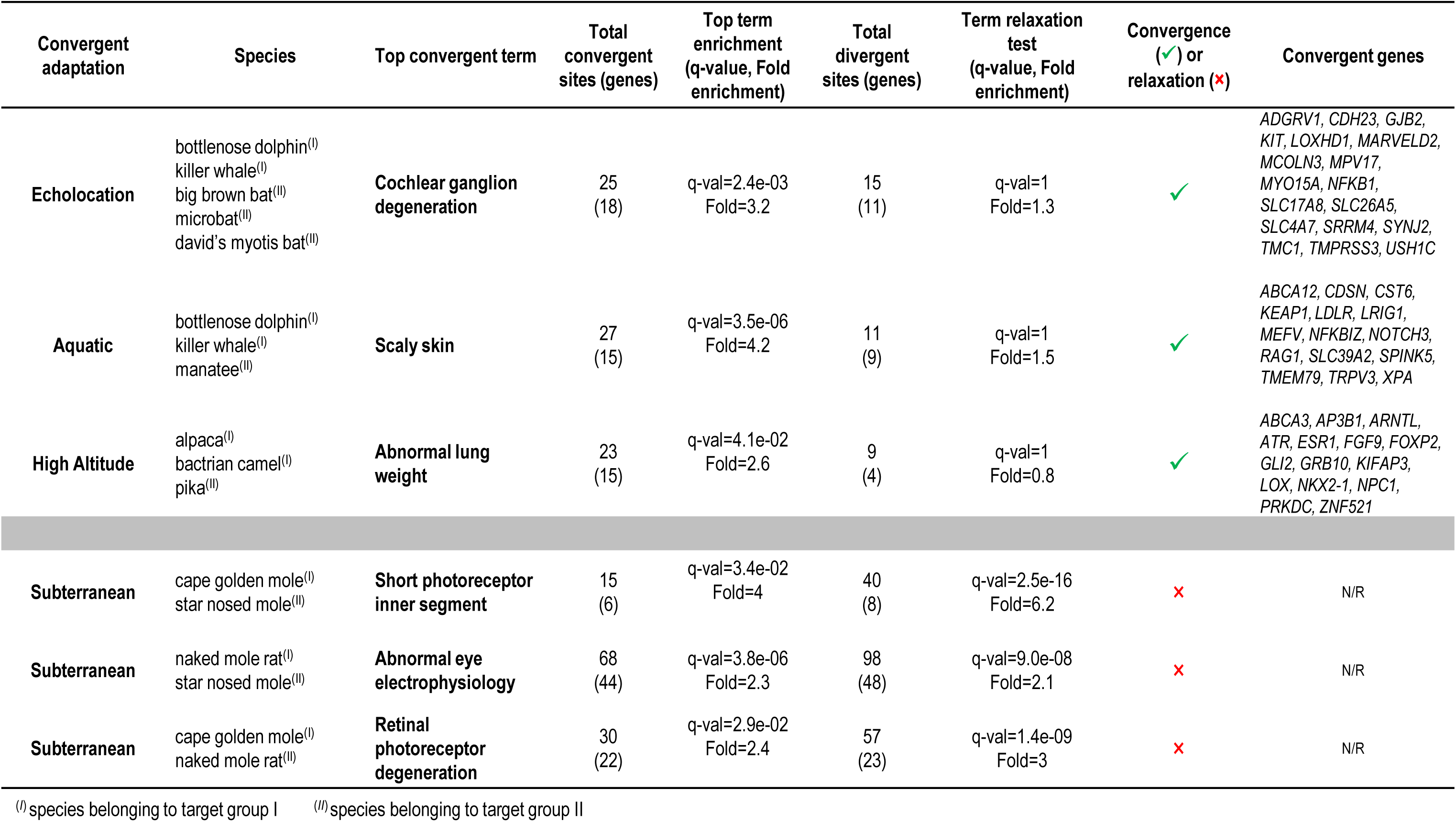
**Finding molecular convergence in convergent phenotypes.** Every row tests a different phenotypic convergence. To declare molecular convergence, the top enriched MGI phenotype term (of 4,000+ tested) for convergent mutations must *not* also be enriched for divergent mutations. We predict convergent molecular evolution in cochlear, skin and lung genes, in echolocating, aquatic and high altitude mammals, respectively, while correctly identifying convergent mutation accumulation in subterranean mammals’ vision genes as resulting from molecular relaxation.

### Making the convergent enrichment test robust to independent relaxation using subterranean mammals

We suspected that independent loss of phenotypic function (vestigialization) can produce a false molecular convergence-like pattern, because relaxed positions of relevant proteins may mutate more freely, and are thus more likely to introduce non-adaptive convergent changes. We sought to test this hypothesis using subterranean mammals whose visual system and multiple associated genes are thought to undergo regressive evolution^20,21^. We selected three independently evolved subterranean mammals: star-nosed mole, naked mole-rat and cape golden mole along with three respective outgroups (Fig. 1a; Supplementary Figure 1). Repeating the analysis described above (and in Fig. 2a), we performed three pairwise convergence tests, between each pair of subterranean mammals (see Methods). The three pairs yielded very similar results: 1024–1250 convergent amino acids in 853–995 MGI annotated genes (Supplementary Tables 2, 4-6), and a top enriched term (*q-value*<0.05 in all cases) directly related to vision: **‘short photoreceptor inner segment’, ‘abnormal eye electrophysiology’** and **‘retinal photoreceptor degeneration’** (Table 1).

Next we hypothesized that when the signal comes from a relaxation event (but not from a convergence event), the same set of genes will also experience a significant number of divergent mutations of conserved amino acid positions (Fig. 1c). Scanning conserved amino acids for divergence events exactly on the branch separating target groups from their outgroups, where both target groups changed the position into a different amino acid, we collected all amino acid divergence events from the three pairs of ‘moles’ (see Methods and Fig. 1c). A total of 1599—2160 divergent amino acids were found in 1147–1391 MGI annotated genes (Supplementary Table 2). When repeating the amino acid enrichment test (Fig. 2a) with the divergent positions, all three top convergent vision terms were found highly significant for divergence (*q-value*<1e-7 in all cases). In contrast, performing the divergence test on the echolocation targets and outgroups, which results in 1131 divergent amino acids in 905 genes (Supplementary Table 2), suggests no enriched accumulation (*q-value*=1) for the top convergent term **‘cochlear ganglion degeneration’.** We conclude by accepting the echolocation signal as resulting from convergent evolution, and rejecting the mole vision signals as likely stemming from independent relaxation (Table 1). We incorporate the relaxation test as an integral part of our test in the remainder.

### Prominent adaptive convergence in skin-related genes of aquatic mammals

We employed the combined convergence and relaxation test to explore the molecular basis for convergent adaptations in aquatic mammals. We scanned for convergent and divergent mutations (801 and 902, respectively; Supplementary Table 2) between toothed whales (bottlenose dolphin, killer whale) and manatee (Fig. 1a; Supplementary Figure 1 and Supplementary Table 7). The most enriched convergent function **‘scaly skin’** (q-value=3.51e-6, 27 amino acids in 15 genes) is not enriched for divergence (*q-value*=1; Table 1). Indeed marine mammals have adapted to aquatic movement drag by reducing pelage hair, and in some cases even frequently replacing the surface layer of their skin^22^.

### Convergent adaptation in high altitude mammals is promoted by coding convergence in lung genes

Finally, we employed our combined test to high altitude adaptation in mammals. We defined the alpaca and Bactrian camel as one target group and pika as the second target (Fig 1a; Supplementary Figure 1); all three species are well-adapted to altitudes higher than 3,000 feet above sea level^23–26^. Our screen revealed 919 convergent and 1220 divergent mutations (Supplementary Table 2). The function most enriched with convergent mutations is **‘abnormal lung weight’** (q-value=0.041, 23 amino acids in 15 genes), which is not enriched with divergent mutations (*q-value*=1; Table 1), suggesting adaptive convergent amino acid substitutions in the respiratory system of the target species. Lung anatomy adaptations to significantly reduced oxygen intake in high-altitude are an active area of research^27^. Interestingly, humans and rodents born at low-oxygen conditions or high altitudes develop larger lungs and increased lung capacities^27,28^.

## Discussion

Convergent phenotypic evolution is a striking and often observed phenomena. The extent to which phenotypic convergence is driven by molecular convergence is a fascinating question with implications on our understanding of the constraints and predictability of species evolution. A second large debate that looms over our work concerns the relative contribution of coding sequence mutations to species evolution^28^. With respect to both of these big questions we show three independent cases where distantly related echolocating, aquatic, and high-altitude mammals display clear signatures of excess accumulation of multiple convergent mutations each, in highly conserved (and thus functionally important) amino acids of genes directly implicated in their phenotypic adaptations.

By their nature, life-style adaptations like the ones we screen here are complex multitissue affairs. The function-specific sets of coding mutations we discover shed a fascinating ray of light onto the underlying genetics of these adaptations. In the context of echolocation, we find multiple mutations previously noted, and discover some new ones. We see the asparagine to threonine (N7T) substitution in *Prestin*, which is essential for a voltage-dependent physiology unique to echolocators^4^, and previously described convergence in *CDH23*^7^ and *TMC1*^29^. We also discover novel convergent events in additional cochlear innervation genes, whose inclusion in our top enriched set strongly suggests important roles in echolocation. Most importantly, we show, in light of ongoing debate^11,12^, that the functional signature of the cochlear ganglion is both statistically strong – the very top of 4,300 different gene sets tested - and absent when shuffling echolocators and outgroups.

Our screen implicitly leverages the fact that (molecular) evolution and (genetic) disease are but two sides of the same coin (the mutation-selection process). The gene function ontology using which we observe our molecular adaptations is rooted in gene mutation phenotypes. For example, the 18 genes we flag in the context of echolocation, have all been observed to cause cochlear ganglion degeneration when mutated. As expected, several of these genes have also been implicated in human hearing loss^30,31^. A nice amino acid level example of “right” and “wrong” mutations (in the right context, at the right time) is the convergent bat & whale F115Y mutation in *GJB2*, found at the center of three codons, each of which is known to independently cause hearing loss in humans when mutated^32^ (Fig. 3a).

**Figure 3.**
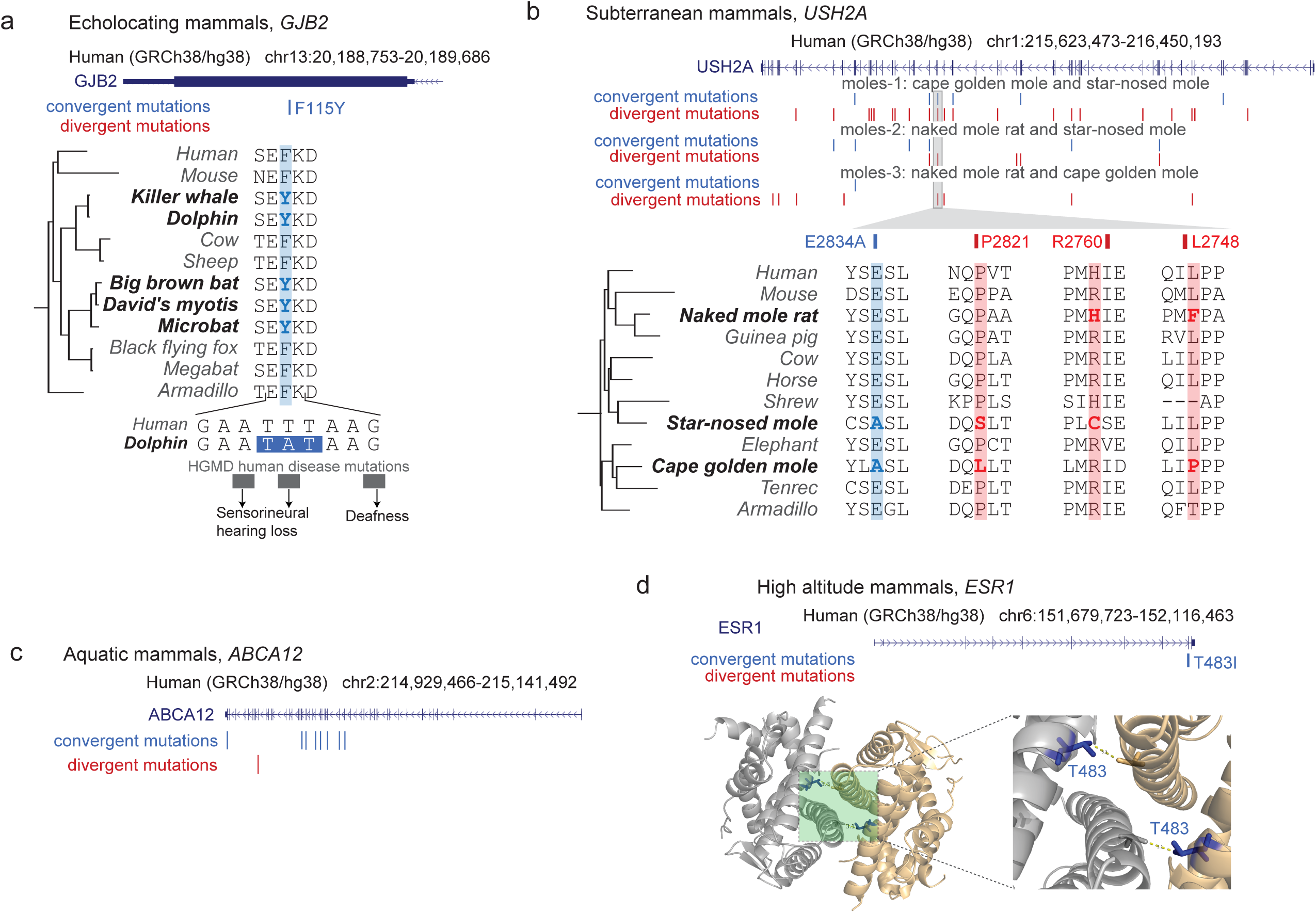
Example convergent and divergent mutations observed. **a,** A convergent mutation F115Y in *GJB2* observed in echolocating mammals is central to three codons containing hearing loss human disease mutations, suggesting the residue’s importance in modulating hearing. **b**, The vision gene *USH2A* contains multiple divergent and convergent mutations in three tested pairs of moles. These mutations likely accumulated as a result of relaxed purifying selection. **c**, Compare to the skin development gene *ABCA12*, which exhibits 8 convergent mutations and only a single divergent mutation in aquatic mammals. **d**, Estrogen receptor gene *ESR1* contains the convergent mutation T483I in high-altitude mammals. Three-dimensional structure of *ESR1* highlights the convergent mutation at the homodimeric interface, suggesting an important functional role.

When applied to aquatic mammals, we find a strong signature for skin adaptation that points to multiple skin development genes. For example, we see no less than 8 convergent mutations in *ABCA12* (Fig. 3c), a skin keratinization gene implicated in Ichthyosis^33,34^. We also identify multiple convergent mutations in TRPV3^10^, a skin gene with roles in themoregulation^35^. In high altitude mammals our screen points to respiratory genes, such as the Q44H mutation in *LOX* (lysil oxydase), a gene with a differential expression pattern in high-altitude chickens^36^ and itself a direct target of the *EPAS1 (HIF2A)* gene proposed to have a role in high-altitude adaptation of Tibetans^37^. We also find a convergent mutation T483I, strikingly located at the protein-protein interface of the estrogen receptor gene *ESR1* (Fig. 3d). Interestingly, estrogen has been proposed to provide a protective effect against hypoxia in women, as high-altitude-induced pulmonary hypertension disproportionately affects men^38^.

As is the case in multiple different evolutionary scenarios, when a surprising number of constrained amino acids rapidly mutate, one must carefully weigh a switch from purifying selection to either positive selection or neutral drift. The subterranean mammals provide an excellent opportunity to fortify our test against the latter scenario. Whereas all three combination of “moles” provide a statistically enriched top term showing an accumulation of convergent mutations in vision genes, all three sets also provide a much stronger statistical signature for accumulated divergent mutations. This is easy to see in *USH2A* where divergent mutations outnumber convergent ones (Fig. 3b) and additional genes on our lists like *CRB1, ABCA3, GUCY2F* and *PDE6C* that have been previously flagged for molecular relaxation in these species^14,21^.

With so many genomes being sequenced, from big centers to individual benchtops, the underpinnings of numerous fascinating phenotypic adaptations are suddenly within reach. To deploy our test one simply needs a set of related sequenced genomes, a gene set functionally annotated in just one of these species, and knowledge of one or more phenotypic convergences of interest among the spanned set of species. Some of the most prominent species clades that fulfil these requirements include yeast, fly, worm, fish and birds.

How does one begin addressing a complex phenotypic adaptation? How can one pick molecular targets which are likely to provide meaningful phenotypes in the lab? Our test provides an attractive first line of attack. Starting from the entire exome, roughly 1-2% of the 3Gb mammalian genome, we first highlight 2.5-3Mb of highly conserved amino acids that are more likely to produce a phenotype when mutated. The stringency of our convergence test lets us detect less than 1,000 amino acid convergence events in each scenario. And finally our most enriched sets offer 23-27 unique amino acid mutations in 15-18 genes, predicted to confer an adaptive advantage in a single relatively well defined function. With wonderful new tools like CRISPR-Cas, and excitingly manipulable systems like inner ear organoids containing functional hair cells^39^, and a three-dimensional model of the lung^40^, both derived from (genetically manipulable) pluripotent stem cells, the time has never been more ripe to use screens like ours to find these fascinating “beachheads” for the exploration of convergent phenotypic evolution, and the contribution of coding sequence mutation to species adaptation.

## Methods

### Species set and gene set

In this study we used genome assemblies of 57 species (listed in Supplementary Table 1), and their substitutions per site weighted phylogenetic tree (Supplementary Figure 1) from UCSC (genome.ucsc.edu). We used human genome assembly GRCh38/hg38 and ENSEMBL (ensembl.org) release 86 human protein-coding gene set as reference.

### Finding conserved, functionally annotated, amino acid positions

We started by picking a pair of independent clades (with one or multiple species) to serve as target groups for our convergent evolution test, and their associated outgroups. The six sets used in this paper are shown in Supplementary Figure 1. We mapped all human genes, except a small fraction overlapping segmental duplication regions (from UCSC), to each of the other 56 species using UCSC liftOver chains^41^. We then excluded genes that were not mapped to at least one species from each of the selected target and outgroups, as well as genes lacking any functional annotations in the Mouse Phenotype Ontology (MGI, more details below). In addition, we required that a gene is aligned in at least 40 of the 56 target species.

We then used the alignments to determine the orthologous amino acid in each available species. Because exon boundaries sometimes shift for evolutionary or for alignment reasons, we excluded the first and last two amino acid positions in each exon from all downstream analyses. We then derived a set of testable amino acids from all remaining genes, where each amino acid is aligned in at least one species from each of the target and outgroups, and is also conserved across all aligned species with Bayesian Branch Length Score>0.9 (see below). We map only the remaining genes back to the MGI ontology, and remove ontology terms annotated by too few or too many genes (see below), and then excluded all genes and amino acids lacking functional annotations in the remaining set of ontology terms. The final sets of ontology terms, genes and amino acids used for testing each of the six species sets in the paper are described in Supplementary Table 2.

### Cross species amino acid conservation score

Using cross-species conservation as a hallmark of functional importance, we required that each amino acid position be aligned in at least 40 placental mammals. We then computed the total substitution per site branch length (BL) over which the dominant amino acid is conserved. This we converted to a Bayesian Branch Length Score (BBLS) which further takes into account the phylogenetic relatedness between species. BBLS was previously demonstrated to outperform BL scores^42^ and is extensively discussed in Xie et al.^43^. We tested only amino-acid positions conserved at a threshold of BBLS>0.9.

### Mouse Phenotype ontology

The MGI Phenotype Ontology catalogs spontaneous, induced, and genetically-engineered mouse mutations and their associated phenotypes^18^. Ontology data (containing 8,949 phenotypic terms; format-version 1.2, May 2016) was lifted over from mouse to human, resulting in 609,253 (canonicalized) gene-phenotype associations. To increase statistical power (by reducing the required multiple testing correction factor in our tests below) we ignored general ontology terms (those annotating more than 500 genes with at least one conserved position) or too specific ontology terms (those annotating fewer than 10 such genes).

### Calling convergent and divergent mutations

Starting from an amino acid alignment at any tested position (derived above) we used PAML (V4.8) to infer the most likely amino acid identity in the internal nodes of our 57 species eutherian phylogenetic tree (Supplementary Figure 1). We scanned only for mutations occurring along the branches leading from the last common ancestor of each target group and its outgroup to the target group itself.

For convergence (Fig. 1b) we called both parallel mutations (i.e., identical mutations in the two independent lineages derived from the same ancestral amino acid) and strictly convergent mutations (identical target mutations derived from different ancestral amino acids) as convergent mutations. We required that the exact same amino acid (the convergent amino acid) is present in each species from both target groups, and that no other amino acids are present in target group species. In target or outgroups containing two or more species (e.g., in the three echolocating bats) we allowed a missing amino acid alignment in some species, as long as the convergent amino acid is represented at least once and no other amino acids are present in the target group.

For divergence (Fig. 1c) we used similar logic: A pair of mutations resulting in different amino acids in the two target groups was called a divergent mutation. In groups of more than one species, we allowed missing alignments (in either target or outgroup) but also allowed different amino acids within target groups, as long as the criteria of amino acid mutation along the branch from the common ancestor with the outgroup to the target itself is satisfied.

### Enrichment test over amino acids

Different genes contribute to our test a different number of conserved (BBLS>0.9) amino acids, depending on gene length and its cross-species conservation. Therefore, rather than test ontology term enrichment over genes, we tested over the conserved amino acid positions, associating each gene’s ontology annotations to all of its amino acid positions we intended to test for convergence/divergence events. This is equivalent to a gene based test where each gene is weighted by the number of positions tested in it.

For a given pair of target lineages, we started by determining the background set of conserved and MGI annotated amino acids as above (Supplementary Table 2). We used the hypergeometric test to calculate functional enrichment over this set. More specifically, the hypergeometric test was executed separately for each ontology term (*π*) with four parameters: (i) *N*, the total number of conserved and annotated amino acid positions (background); (ii) K_*π*_, the total number of positions annotated by an ontology term (a subset of the background); (iii) n, the total number of positions with convergent (or divergent) mutations identified from the background; and (iv) k_*π*_, the intersection between ii and iii. We computed a standard hypergeometric p-value of the observed enrichment for term *π* as the fraction of ways to choose n (converged or diverged) amino acid positions without replacement from the entire group of N positions such that at least k_*π*_ of the n have ontology annotation *π*, using the formula below:

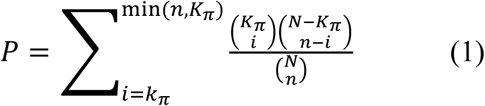

We corrected all p-values for multiple testing by setting a Benjamini-Hochberg False Discovery Rate (FDR) of 0.05 for significance. We also computed the fold enrichment for convergent (or divergent) mutations labeled by a given term by dividing the observed fraction of convergent (or divergent) mutations intersecting the tested term by the expected fraction of mutations: (*k_π_* /n)/(*K_π_* /N). We ranked all terms by q-value (corrected p-values), and asked whether the top term passed the threshold for convergence (or divergence) at a False discovery Rate (FDR) lower than 0.05, and Fold enrichment greater than 2.

## Acknowledgements

We thank the members of the Bejerano Laboratory, particularly M. J. Berger, W. Heavner, H. M. Moots, J. H. Notwell, J. Birgmeier, K. A. Jagadeesh, B. Yoo, H. Guturu and A. M. Wenger for technical advice and helpful discussions. We thank H. Clawson (UCSC) for assistance with mammalian genome alignments, and the community at large for the availability of all the different genomes. This work was funded in part by NIH grants U01MH105949 and R01HG008742, a Packard Foundation Fellowship and a Microsoft Faculty Fellowship (GB).

## Author contributions

A.M. and G.B. designed the study, analyzed results and wrote the manuscript. Y.T., M.G. and B.A.B provided computational tools. M.G. and H.I.C helped with the analysis. All authors commented on the manuscript.

## Supplementary Information legend

**SI Table 1**

**Species list.** The list of 57 placental mammal species and genome assemblies used.

**SI Table 2**

**Key statistics from our molecular convergence test.** For each of the six scenarios tested we show: The number of MGI terms tested for convergence and divergence, the total number of amino acid positions tested and the number of genes containing them, the number of convergent (divergent) mutations detected, and the number of genes harboring them. Moles-1: cape golden mole and star-nosed mole; moles-2: naked mole rat and star-nosed mole; moles-3: cape golden mole and naked mole rat.

**SI Tables 3-8**

**All observed convergent and divergent mutations.** A full list of MGI annotated convergent and divergent mutations and associated genes identified in each test at conserved amino-acid positions (see Methods). Genomic coordinates for the convergent positions are given relative to the reference genome (human, GRCh38/hg38). SI Table 3) echolocation, SI Table 4) subterranean mammals, moles-1: cape golden mole and star-nosed mole, SI Table 5) subterranean mammals, moles-2: naked mole rat and star-nosed mole, SI Table 6) subterranean mammals, moles-3: naked mole rat and cape golden mole, SI Table 7) aquatic mammals, SI Table 8) high-altitude mammals.

**SI Figure 1.**
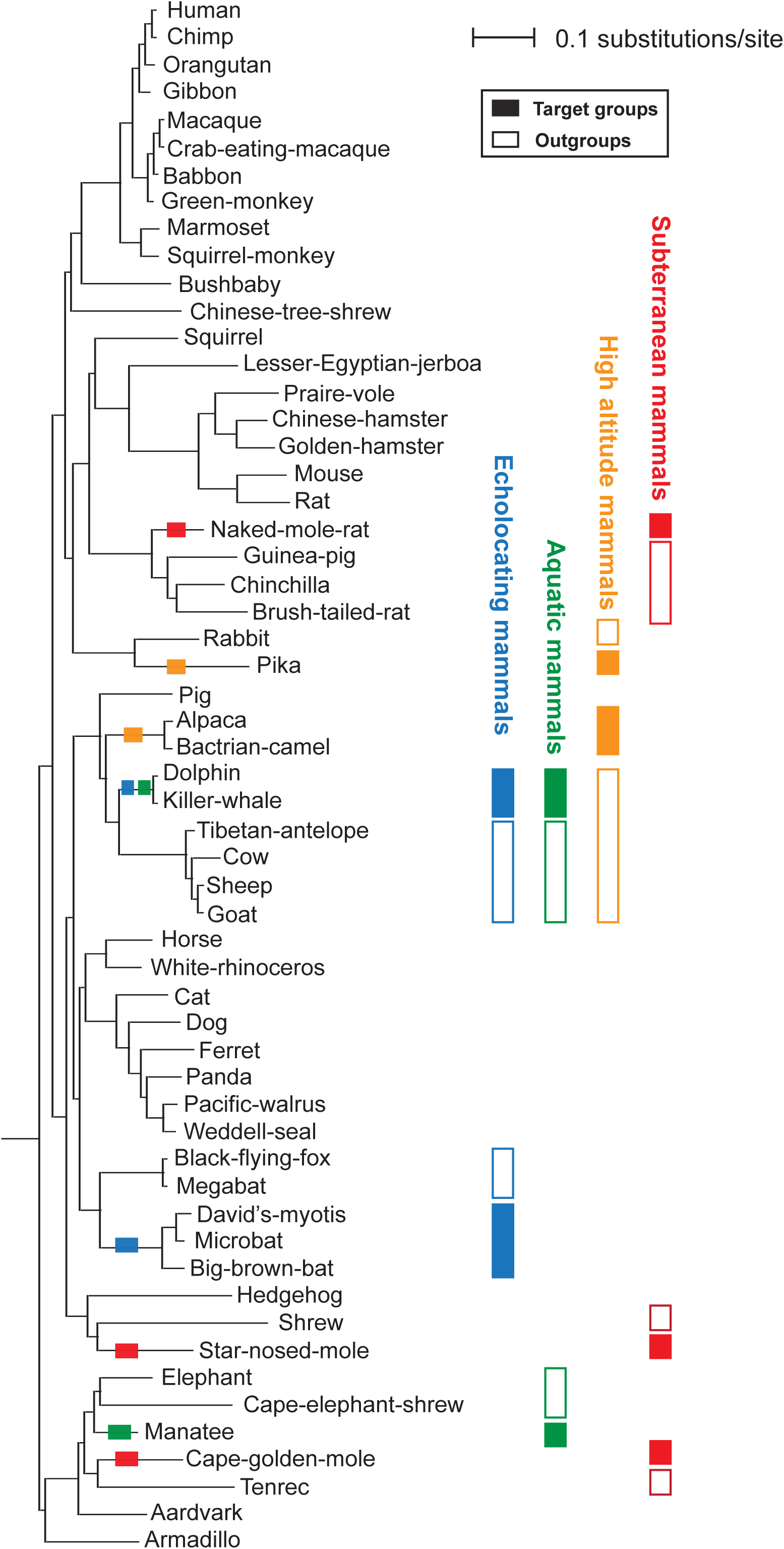
Weighted phylogenetic tree. A substitution per site weighted tree of the 57 species used in our screen. Filled rectangles mark target groups and empty rectangles mark outgroups for lineages with independent phenotypic evolution of echolocation (blue), aquatic (green), high-altitude (orange) and subterranean (red) lifestyles.

